# Vitamin C triggers NF-κB-driven epigenomic reprogramming and enhanced immunogenic responses of dendritic cells

**DOI:** 10.1101/2022.05.26.493381

**Authors:** Octavio Morante-Palacios, Gerard Godoy-Tena, Josep Calafell-Segura, Laura Ciudad, Eva M. Martínez-Cáceres, José Luis Sardina, Esteban Ballestar

**Author notes:** Correspondence: E. Ballestar.

## Abstract

Dendritic cells (DCs) are central in the immune system, bridging the adaptive and innate immune responses. Research on *in vitro* differentiation of DCs from monocytes provides both in-depth understanding of the analogous in vivo process and potential sources for cancer cell therapy. Active DNA demethylation is crucial in DC differentiation. Vitamin C is a known cofactor of ten-eleven translocation (TET) enzymes, which drive active demethylation. Currently, the effects of vitamin C treatment on human immune cells are poorly understood. In this study, we have studied the epigenomic and transcriptomic reprogramming orchestrated by vitamin C in monocyte-derived DC differentiation and maturation. Vitamin C triggers extensive demethylation at NF-kB/p65 binding sites, together with concordant upregulation of antigen-presentation immune response-related genes during DC maturation. p65 interacts with TET2 and mediates the aforementioned vitamin C-mediated changes, as demonstrated by pharmacological inhibition. Moreover, vitamin C increases TNFβ production in DCs through NF-kB, in concordance with the upregulation of its coding gene and the demethylation of adjacent CpGs. Finally, vitamin C enhances DC’s ability to stimulate the proliferation of autologous antigen-specific T cells. We propose that vitamin C can improve monocyte-derived DC-based cell therapies. Finally, our results provide a feasible mechanism of action for intravenous high-dose vitamin C treatment in patients.

## INTRODUCTION

Dendritic cells (DCs) play a central role in the immune system, bridging innate and adaptive immune responses. As innate immune cells, they are able to recognize a plethora of pathogen-associated molecular patterns (PAMPs) and damage-associated molecular patterns (DAMPs) through pattern recognition receptors (PRRs)(*1*). Moreover, they are very efficient at antigen processing and presentation to T cells and are, therefore, responsible for initiating antigen-specific immune responses.

DCs are very heterogeneous, comprising plasmacytoid DCs (pDCs) and conventional DCs (cDCs). Furthermore, monocyte-derived DCs (moDCs) can be differentiated *in vitro* from monocytes (MOs), with GM-CSF and IL-4(*2*). moDCs have been classically used as a convenient model that mimics blood DCs, especially cDC2(*3*). However, increasing evidence indicates that MOs can extravasate to peripheral tissues and give rise to *bona fide* moDCs *in vivo*(*4*).

Changes in DNA methylation, mainly active DNA demethylation, are involved in several differentiation processes from MOs, including macrophages(*5*), osteoclasts(*6*), and DCs(*7*). and are crucial for immune cell differentiation, identity, and function(*8*). In general, DNA demethylation has been observed to be more extensive during MO differentiation than during subsequent maturation/activation(*9*). Additionally, the maturation of moDCs with live bacteria produces DNA demethylation that follows gene activation, limiting the potential direct regulatory effects of DNA methylation in such context(*10*). TET2, a member of the Ten-Eleven Translocation (TET) methylcytosine dioxygenases, is involved in multi-step active demethylation processes, and is critical for terminal MO-related differentiation(*6*, *11*). Recently, TET2 has also been implicated in glucocorticoid- and vitamin D-mediated modulation of the immunogenic properties of DCs(*12*, *13*).

Vitamin C (L-ascorbic acid) is an essential nutrient with pleiotropic functions. Its deficiency is associated with a disease, namely scurvy, characterized by a plethora of symptoms, including the malfunction of the immune system. For instance, the normal intracellular level of vitamin C in MO cytoplasm is ~3mM, 60 times higher than the plasma level, reflecting a specific function of the molecule in immune cell biology. Vitamin C can act as a cofactor of Fe-containing hydroxylases such as TET enzymes and Jumonji C domaincontaining histone demethylases (JHMDs), enhancing their catalytic activity(*14*). Some studies in mice suggest that vitamin C can stimulate DC capacity to produce proinflammatory cytokines and promote differentiation of T cells(*15*). Moreover, vitamin C intravenous treatment in mice has been shown to abrogate cancer progression through direct TET2 function restoration in cancer cells(*16*) and immune system modulation(*17*).

The *in vivo* modulation of DC migration and function, and the administration of DC-based vaccines, are promising strategies to treat different types of cancer(*18*). In particular, autologous moDCs obtained *ex vivo* from patient-derived blood MOs have been used in several clinical trials with mixed results(*19*–*21*). In this regard, the improvement of moDC generation *in vitro*, and the use of molecules to modulate MO differentiation *in vivo* may boost the clinical outcome of cancer patients.

In this work, we have investigated the effects of vitamin C treatment during MO-to-DC *in vitro* differentiation and maturation, identifying extensive DNA demethylation associated with the upregulation of migration, chemotaxis, antigen presentation, and immune response-related genes. Moreover, vitamin C-mediated DNA demethylation, gene upregulation, and increased TNFβ production during DC maturation were associated with p65, a component of the NF-κB complex that interacts with TET2 in this context. We have shown how the modulation of DNA methylation changes during DC differentiation and maturation can yield functional and phenotypic changes in these cells, improving their immunogenicity.

## RESULTS

### Vitamin C drastically enhances DNA demethylation during monocyte to dendritic cell differentiation and maturation

MOs isolated from peripheral blood of healthy donors were differentiated *in vitro* to DCs for 7 days using GM-CSF and IL-4, in the presence or absence of vitamin C (vitC). Samples were collected on day 2, in the middle of the differentiation (Ø / Ø_vitC_), and on day 7, including immature DCs (iDCs) (iDC / iDC_vitC_), without further treatment, and mature DCs, exposed the last 48h to lipopolysaccharide (LPS) (mDC / mDC_vitC_) (Figure 1A).

**Figure 1.**
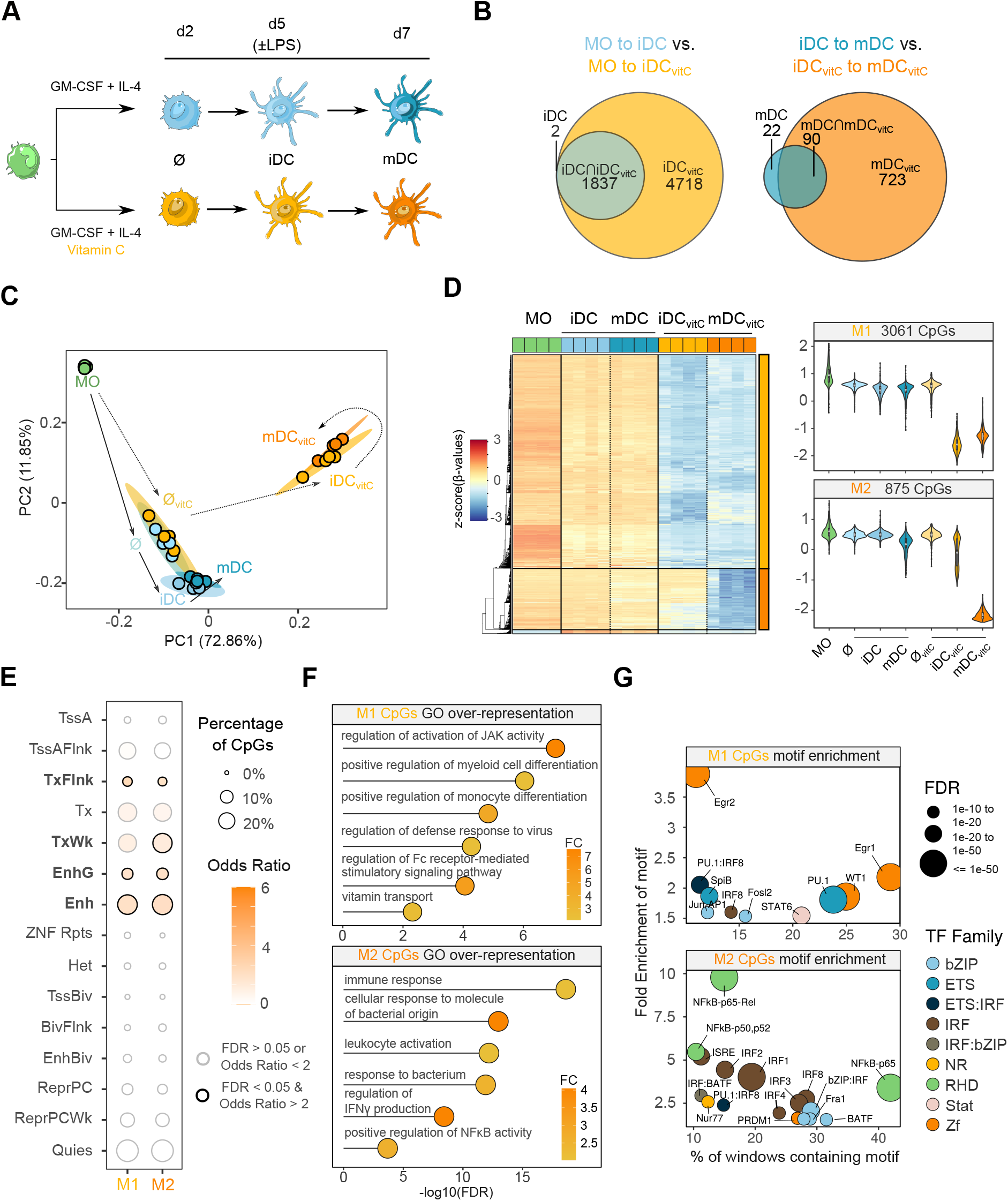
Vitamin C-mediated dendritic cell DNA methylome remodeling. (A) Scheme depicting the experimental setup. Monocytes (MO) were differentiated to dendritic cells (DCs) using GM-CSF and IL-4, in the presence or absence of vitamin C (vitC). Samples were collected on day 2, in the middle of the differentiation, and on day 7, including immature DCs (iDCs) and mature DCs (mDCs), exposed the last 2 days to lipopolysaccharide (LPS). (B) Area-proportional Venn diagrams comparing the demethylated CpG sets of MO-to-iDC vs. MO-to-iDC_vitC_, and iDC-to-mDC vs. iDC_vitC_-to-mDC_vitC_ transitions. (C) Principal Component Analysis (PCA) of differentially methylated CpGs comparing all groups pairwise. Principal component 1 and principal component 2 are represented on the x- and y-axis, respectively. (D) DNA methylation heatmap of differentially methylated CpGs comparing iDC to iDC_vitC_, and mDC to mDC_vitC_ (Δβ ≥ 0.3, FDR < 0.05). Scaled β-values are shown (lower DNA methylation levels in blue and higher methylation levels in red). On the right side, violin plots of clusters M1 and M2 depict scaled β-values. (E) Enrichment of M1 and M2 CpGs in ChromHMM 15-states categories of MOs (Roadmap Epigenomics Project). Significantly enriched categories (FDR < 0.05 and odds ratio > 2) are depicted with a black stroke, including TxFlnk (Flanking Active TSS), TxWk (Weak Transcription), EnhG (Genic Enhancers), and Enh (Enhancers). (F) GO (Gene Ontology) over-represented categories in M1 and M2 CpGs. Fold Change in comparison with background (EPIC array CpGs) and −log10(FDR) is represented. (G) Bubble scatterplot of transcription factor binding motif enrichment for M1 and M2 CpGs. The x-axis shows the percentage of windows containing the motif and the y-axis shows the fold enrichment of the motif over the EPIC background. Bubbles are colored according to the transcription factor family. FDR is indicated by bubble size.

DNA methylation was profiled using Illumina Infinium MethylEPIC arrays, which cover around 850,000 positions in the human genome. First, overall changes in DNA methylation were calculated between groups pairwise(Supplementary Figure 1A). DNA demethylation was the most prominent change, as previously described(*7*). We then compared the demethylated positions in MO-to-iDC differentiation, in comparison with MO-to-iDC_vitC_ differentiation, as well as iDC-to-mDC maturation in comparison with iDC_vitC_-to-mDC_vitC_ maturation. As it can be observed, the positions demethylated in the differentiation and maturation processes in the presence of vitamin C include the majority of CpGs demethylated in the regular differentiation and maturation of DCs, as well as a vast number of additional CpGs(Figure 1B).

Principal component analysis (PCA) revealed that on day 2, most DNA methylation variance of the MO-to-iDC differentiation had already developed (Figure 1C), whereas no differences with the vitamin C stimulus were found. In contrast, on day 7, vast differences were observed for both iDC_vitC_ and mDC_vitC_, in comparison with their corresponding controls without vitamin C suggesting that the vitamin C-mediated boost in demethylation occurs later in time. The variable that explains most of the variance in DNA methylation resides in the presence/absence of vitamin C during differentiation.

All differentially methylated positions (DMPs) associated with vitamin C (iDC vs. iDC_vitC_ and mDC vs. mDC_vitC_) were represented together, revealing two clusters of CpG sites (M1 and M2) (Figure 1D, Supplementary Table 1). M1 corresponds with CpGs demethylated during differentiation in the presence of vitamin C whereas M2 are CpGs demethylated during LPS-mediated maturation in the presence of vitamin C. Both clusters were enriched in monocytic enhancers and regions flanking active transcription start sites (Figure 1E). That corresponds with predominant localization in intergenic regions, far from CpG islands (Supplementary Figure 1B, Supplementary Figure 1C). Of note, M1 and M2 CpGs are located in regions with subtle increases in H3K27ac and H3K4me1 histone marks from MOs to iDCs and mDCs, respectively. This suggest that these regions are primed for activation, even in the absence of vitamin C (Supplementary Figure 1D).

Gene Ontology enrichment analysis of M1 CpGs-associated genes revealed categories related to positive regulation of myeloid differentiation, regulation of JAK activation, regulation of defense response to virus, and vitamin transport, among others. In contrast, M2 CpGs were enriched in terms related to LPS response and immune activation such as cellular response to molecules of bacterial origin, leukocyte activation, response to bacterium, regulation of IFNγ production, and positive regulation of NF-κB activity (Figure 1F). These functions are consistent with the respective association of M1 and M2 clusters with the DC differentiation and maturation steps.

Active demethylation in myeloid cells is often mediated by transcription factors that recruit specific epigenetic enzymes. In this regard, M1 CpGs were enriched in the consensus binding motifs of transcription factors previously related to DC differentiation, such as EGR2(*22*), STAT6(*7*), and PU.1(*22*), whereas M2 CpGs were enriched in the consensus binding motifs of NF-kB, AP-1, and IRF (Figure 1G).

### Vitamin C drives gene expression remodeling in dendritic cells

Given the extensive differences in DNA methylation mediated by vitamin C, we then performed RNA-seq of MOs, Ø, Ø_vitC_, iDC, iDC_vitC_, mDC, and mDC_vitC_ and checked for differences in their transcriptomes. In contrast with DNA methylation, transcriptome variance of principal component (PC)1 and PC2 are mainly explained by the maturation of DCs and the differentiation of MO to DC, respectively. However, differences between iDC and iDC_vitC_ and mDC and mDC_vitC_ can also be observed in the PCA (Figure 2A).

**Figure 2.**
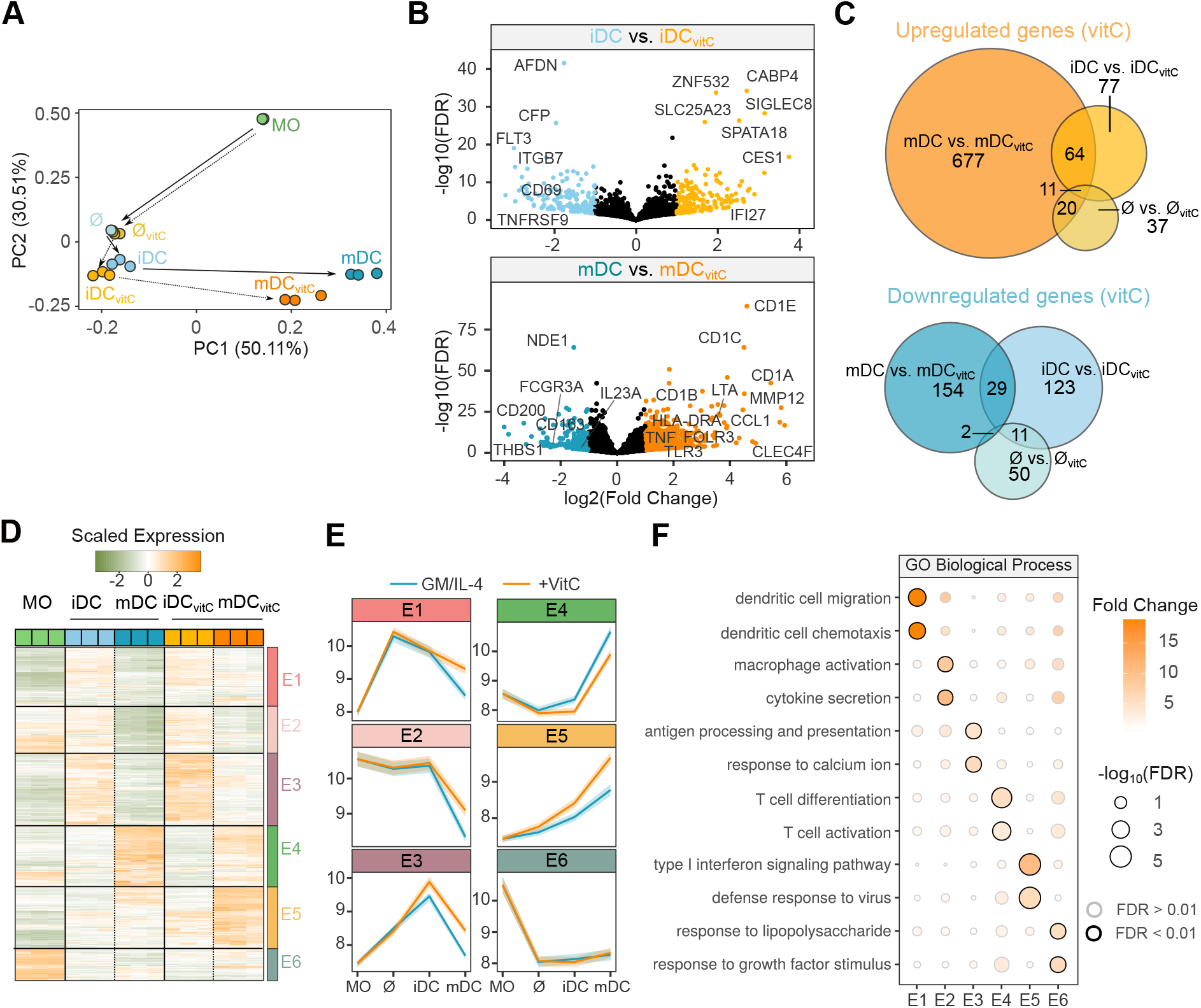
Shifting in gene expression of dendritic cells triggered by vitamin C. (A) Principal Component Analysis (PCA) of gene expression. Principal component 1 and principal component 2 are represented on the x- and y-axis, respectively. (B) Volcano plots of gene expression in the iDC vs. iDC_vitC_ and the mDC vs. mDC_vitC_ comparisons. The binary logarithm of the fold change is represented on the x-axis, whereas the negative decimal logarithm of the FDR is represented on the y-axis. Downregulated genes are shown in blue (FDR < 0.05, Fold Change < −2) and upregulated genes are shown in orange (FDR < 0.05, Fold Change > 2). (C) Area-proportional Venn diagrams comparing the upregulated and downregulated gene sets of the Ø vs. Ø_vitC_, iDC vs. iDC_vitC_, and mDC vs. mDC_vitC_ comparisons. (D) Gene expression heatmap of differentially expressed genes comparing Ø to Ø_vitC_, iDC to iDC_vitC_, and mDC to mDC_vitC_ (absolute logFC > 1, FDR < 0.05). Scaled expression VST values are shown (lower expression levels in green and higher expression values in orange). The division of the heatmap with the ward.D2 clustering method yielded 6 different expression clusters (E1-E6). (E) Temporal progression of gene expression of the expression clusters (E1-E6) during the differentiation process with (orange) or without (blue) vitamin C. The y-axis shows VST values, where a higher value means a higher gene expression, and the line ribbons represent the 95% confidence interval. (F) Gene Ontology (GO) over-representation of GO Biological Process categories in the E1-E6 clusters. Fold Change of genes over the background and −log10(FDR) of the Fisher’s exact tests are shown. Significant categories (FDR < 0.01) are depicted with a black stroke.

Since samples were collected on day 2 (Ø / Ø_vitC_), and on day 7, without (iDC / iDC_vitC_) or with (mDC / mDC_vitC_) activation, three potential comparisons of vitamin C-treated cells can be performed, in relation to their respective controls. On day 2, very few differences were found between Ø and Ø_vitC_ (63 downregulated and 75 upregulated genes). On day 7, we found 163 downregulated and 159 upregulated genes between iDC and iDC_vitC_, whereas most differences were found between mDC and mDC_vitC_ (185 downregulated and 772 upregulated genes) (Figure 2B, Supplementary Table 2). Furthermore, most differentially expressed genes (DEGs) were not shared between comparisons (Figure 2C).

We then joined the differentially expressed genes of the three comparisons and divided them into six clusters (expression clusters E1-E6) with different behaviors (Figure 2D, Supplementary Table 3). E1, E2, and E3 clusters show a diminished downregulation trend in the iDC_vitC_ to mDC_vitC_ transition, in comparison with the iDC to mDC transition, whereas E5 genes are more upregulated in the iDC_vitC_ to mDC_vitC_ transition in comparison with the iDC to mDC transition (Figure 2E).

Gene Ontology Enriched terms were calculated for each expression cluster, obtaining distinctive categories (Figure 2F). For instance, E1 was enriched in dendritic cell migration and chemotaxis, E2 in macrophage activation and cytokine secretion, E3 in antigen processing and presentation and response to calcium ion, and E5 in type I interferon signaling pathway and defense response to virus.

### Vitamin C-mediated demethylation is linked to increased gene expression during dendritic cell maturation

To identify potential functional effects of the vitamin C-mediated demethylation, we linked each DMP with its closest gene. When the global profile of M1 and M2 associated genes was intersected with DEGs from Figure 2D, we found that M1 associated genes are less downregulated in mDCs with vitamin C treatment, whereas M2 associated genes are upregulated in mDCs with vitamin C treatment (Figure 3A). Concordantly, E1 and E2 clusters are enriched in M1-associated genes, and the E5 cluster is enriched in M2-associated genes (Figure 3B).

**Figure 3.**
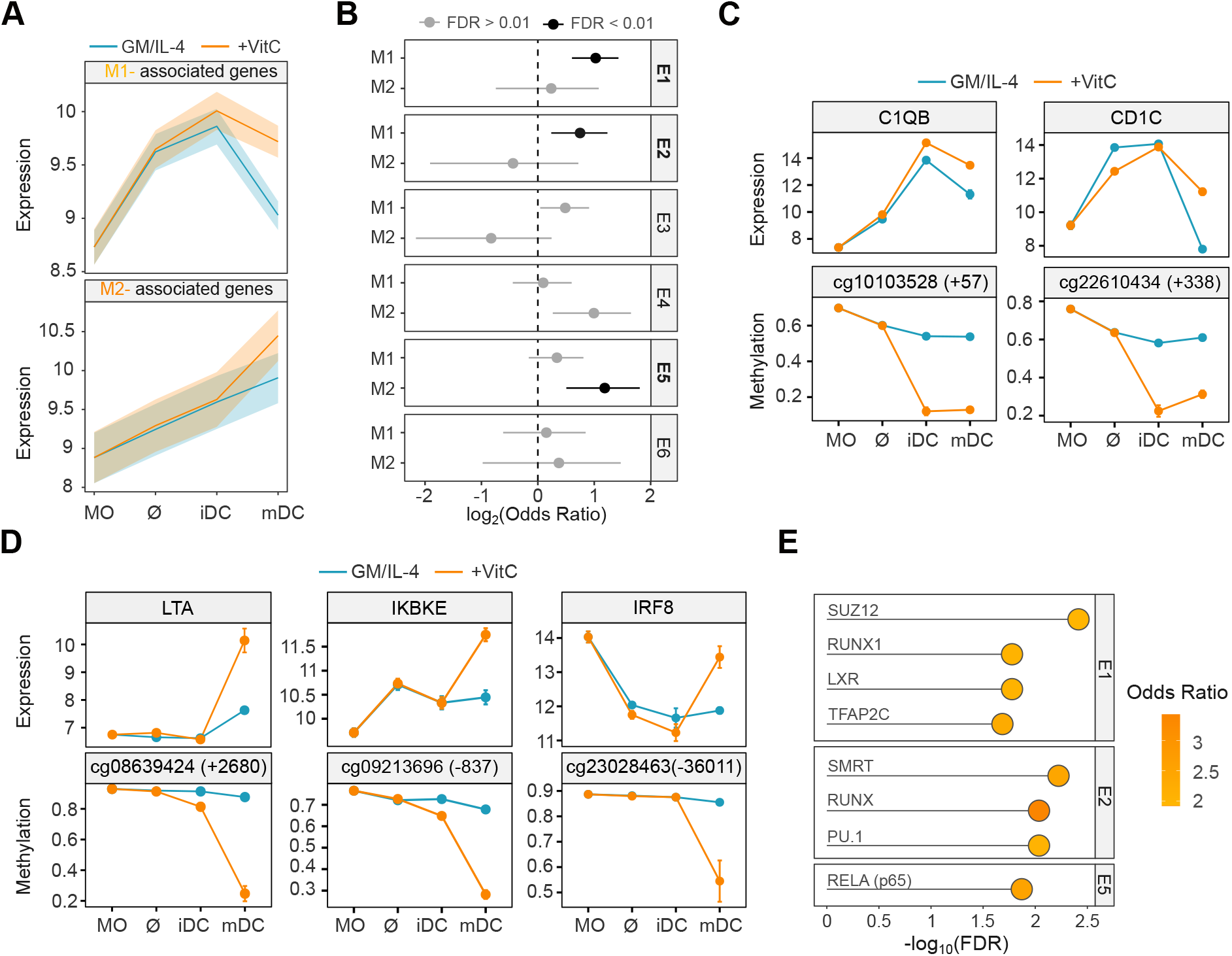
Integration of DNA methylation and gene expression vitamin C-mediated remodeling. (A) M1 and M2 CpGs were associated with the nearest transcription start site. Temporal progression of gene expression of the M1- and M2-associated genes intersected with vitamin C-mediated differentially expressed genes (Figure 2D) with (orange) or without (blue) vitamin C is represented. The y-axis shows VST values, where a higher value means a higher gene expression, and the line ribbons represent the 95% confidence interval. (B) Enrichments of M1- and M2-associated genes over the E1-E6 expression clusters were calculated using Fisher’s exact tests. Odds ratios ± 95% confidence intervals are shown. Significant enrichments (FDR < 0.01) are shown in black. (C) Selected examples of M1 CpGs and associated genes. Temporal progression of DNA methylation (β-value) (below) and gene expression (VST) (above) during the differentiation process with (orange) or without (blue) vitamin C is depicted. (D) Selected examples of M2 CpGs and associated genes. Temporal progression of DNA methylation (β-value) (below) and gene expression (VST) (above) during the differentiation process with (orange) or without (blue) vitamin C is depicted. (E) Enrichment of clusters with gene expression / DNA methylation correlation with genesets from CheA 2016 database, containing genes putatively regulated by transcription factors. The odds ratio over the background and −log10(FDR) of the Fisher’s exact tests are shown.

Individual DMPs and their associated DEGs illustrate different relationships between DNA methylation/gene expression. M1-associated genes, C1QB and CD1C, which are related to the complement system and antigen presentation respectively, show demethylation with vitamin C treatment in iDCs, conjoined with a lower reduction in expression after activation (Figure 3C). Furthermore, M2-associated genes such as LTA, IKBKE, and IRF8 related to TNFβ production, NF-kB pathway regulation, and interferon regulation respectively, depict a decreased methylation in mDC with vitamin C, concomitant with increased gene expression (Figure 3D).

Finally, a database of transcription factor regulation(*23*) was used to infer potential transcription factors involved in regulating expression clusters associated with DNA methylation changes (Figure 3E). Interestingly, PU.1 and RELA (p65) were associated with M1/E2 and M2/E5 clusters, respectively.

### NF-κB/p65 orchestrates vitamin C-mediated DNA demethylation, gene upregulation, and increased proinflammatory cytokine production

Since p65 was associated with both DNA demethylation (Figure 1G) and gene upregulation (Figure 3E) during mDC_vitC_ maturation, we studied the protein expression and phosphorylation by Western Blot. First, we found that p65 presents similar protein levels in iDCs, mDCs, iDC_vitC_, and mDC_vitC_. However, phosphorylated p65 (Ser536) (p-p65) is increased in both mDCs and mDC_vitC_ (Figure 4A). Moreover, we also detected p-p65 in the nuclear fraction (NF) of mDCs and mDC_vitC_, enabling it to act as a transcription factor (Figure 4B).

**Figure 4.**
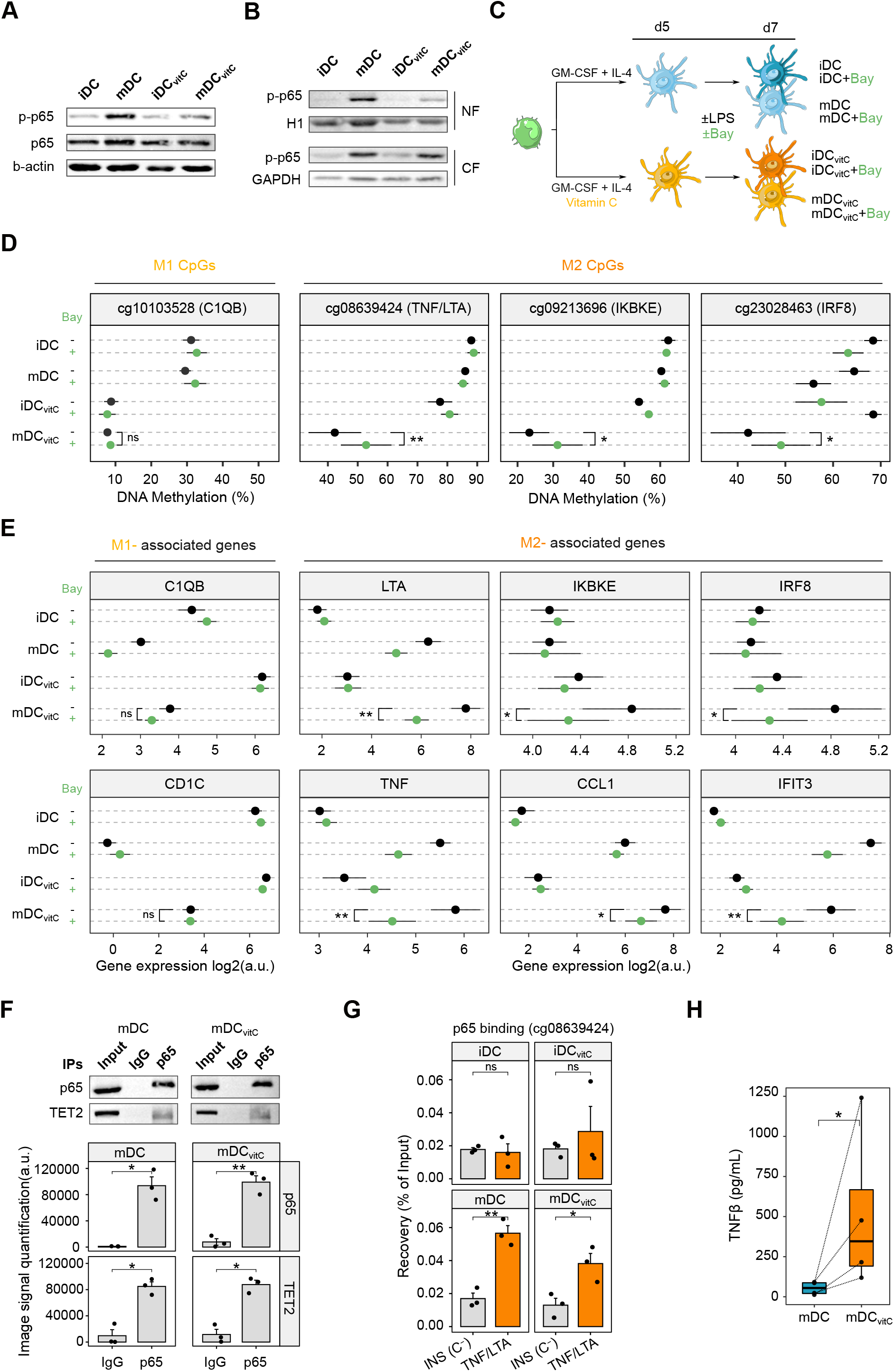
Role of p65 in the transcriptomic and epigenomic remodeling of vitamin C-treated dendritic cells. (A) Western blot of phosphorylated p65 (p-p65) (Ser536) and total p65 in whole-cell lysates. β-actin was used as a loading control. (B) Western blot of p-p65 in the nuclear (NF) and cytoplasmic fractions (CF). GAPDH and histone H1 proteins were used as loading control for cytoplasms and nuclei, respectively. (C) Scheme depicting the inhibition of p65. Monocytes (MO) were differentiated to dendritic cells (DCs) using GM-CSF and IL-4, in the presence or absence of vitamin C (vitC). On day 5, lipopolysaccharide (LPS) and BAY 11-7082 (Bay) or equivalent amounts of diluent were added to the cell culture. On day 7, iDC/iDC_vitC_, iDC + Bay /iDC_vitC_ + Bay, mDC/mDC_vitC_, and mDC + Bay/ mDC_vitC_ + Bay were obtained. (D) DNA methylation of selected CpGs from the M1 and M2 clusters. The average DNA methylation obtained with pyrosequencing is represented with points, and the black lines indicate the standard error of the mean. P-values of paired t-tests are shown (n = 4) (ns P>0.05, *P < 0.05, **P < 0.01). (E) The average gene expression obtained with RT-qPCR is represented with points, and the black lines indicate the standard error of the mean. P-values of paired t-tests are shown (n = 6) (ns P>0.05, *P < 0.05, **P < 0.01). (F) Western blot of the co-immunoprecipitation of p65, showing the signal of p65 and TET2 proteins. Below, the image signal quantifications of three independent western blots are shown for each protein. P-values of paired t-tests are shown (n = 3) (mean ± standard error of the mean) (*P < 0.05, **P < 0.01). (G) Chromatin Immunoprecipitation (ChIP) signal of p65 binding to a negative control locus (*INS*) and around an M2 CpG (cg08639424) associated with *LTA* and *TNF*. The RT-qPCR signal relative to the ChIP input is shown (n = 3). P-values of t-tests are shown (n = 3) (mean ± standard error of the mean) (ns P > 0.05, *P < 0.05). (H) TNFβ production of mDCs and mDC_vitC_, after 5 days of differentiation and 48h of maturation with LPS. The P-value of a Wilcoxon rank-sum test is shown (n=3)

To further explore the role of NF-κB/p65 in the mDC_vitC_ transcriptomic and epigenomic reprogramming, we utilized a chemical inhibitor of IκB degradation (BAY 11-7082, Bay) which reduces the nuclear translocation of p65 (*24*, *25*) (Supplementary Figure 2A). MOs were differentiated in DCs as previously described, adding Bay (10μM) or Dimethyl Sulfoxide (DMSO) on day 5 (Figure 4C).

Firstly, we found that the demethylation of CpGs from the M2 cluster was partially reverted inhibiting NF-kB in mDC_vitC_, concordantly with its enrichment in p65 motifs. Conversely, a CpG from the M1 cluster was not affected (Figure 4D). Secondly, we also tested the expression of M2-associated genes, revealing that, inversely to DNA methylation, gene expression decreases in mDC_vitC_ when NF-κB is inhibited. However, no differences were found in M1-associated genes (Figure 4E).

We then checked the potential interaction between p65 and TET2, a key mediator of active demethylation in myeloid cells whose activity is enhanced by vitamin C. The coimmunoprecipitation of p65 revealed its interaction with TET2 in both mDCs and mDC_vitC_ (Figure 4F).

Additionally, we performed a ChIP-qPCR analysis of p65 on day 6, comparing the binding in a negative control amplicon around the *INS* gene with an amplicon around the cg08639424, located close to *LTA* and *TNF* genes. p65 was significantly enriched in mDCs/mDC_vitC_, but not in iDCs/iDC_vitC_ (Figure 4G).

Since the *LTA* gene, which encodes TNFβ, was found upregulated and adjacent CpGs become demethylated in mDC_vitC_, we measured the TNFβ protein levels secreted by these cells in comparison with mDCs. mDCs produced little amounts of TNFβ whereas mDC_vitC_ supernatant contained considerably higher concentrations (Figure 4H). Moreover, Bay treatment damped TNFβ production of mDC_vitC_, consistent with the reversion in the cg08639424 demethylation and LTA upregulation (Supplementary Figure 2B).

### Vitamin C produces dendritic cells with higher T cell stimulation capabilities

We then characterized the mDC_vitC_ phenotype in comparison with mDCs. In particular, we studied the T cell stimulation capabilities and antigen presentation of mDC_vitC_ in contrast to mDCs. First, we performed a clonal expansion of T cells from healthy donors with a SARS-CoV-2 mix of antigens. After the process, we verified that 95% of the resulting cells were T lymphocytes by staining them with anti-CD4 and anti-CD8 antibodies (Supplementary Figure 3A, Supplementary Figure 3B). We then differentiated autologous MOs *in vitro* to iDC_vitC_/iDCs. After 48h of maturation with LPS and 1h of antigen loading with the same SARS-CoV-2 antigen mix, mDC_vitC_/mDCs were cocultured with Carboxyfluorescein succinimidyl ester (CFSE)-stained clonal T cells for 5 days (Figure 5A).

**Figure 5.**
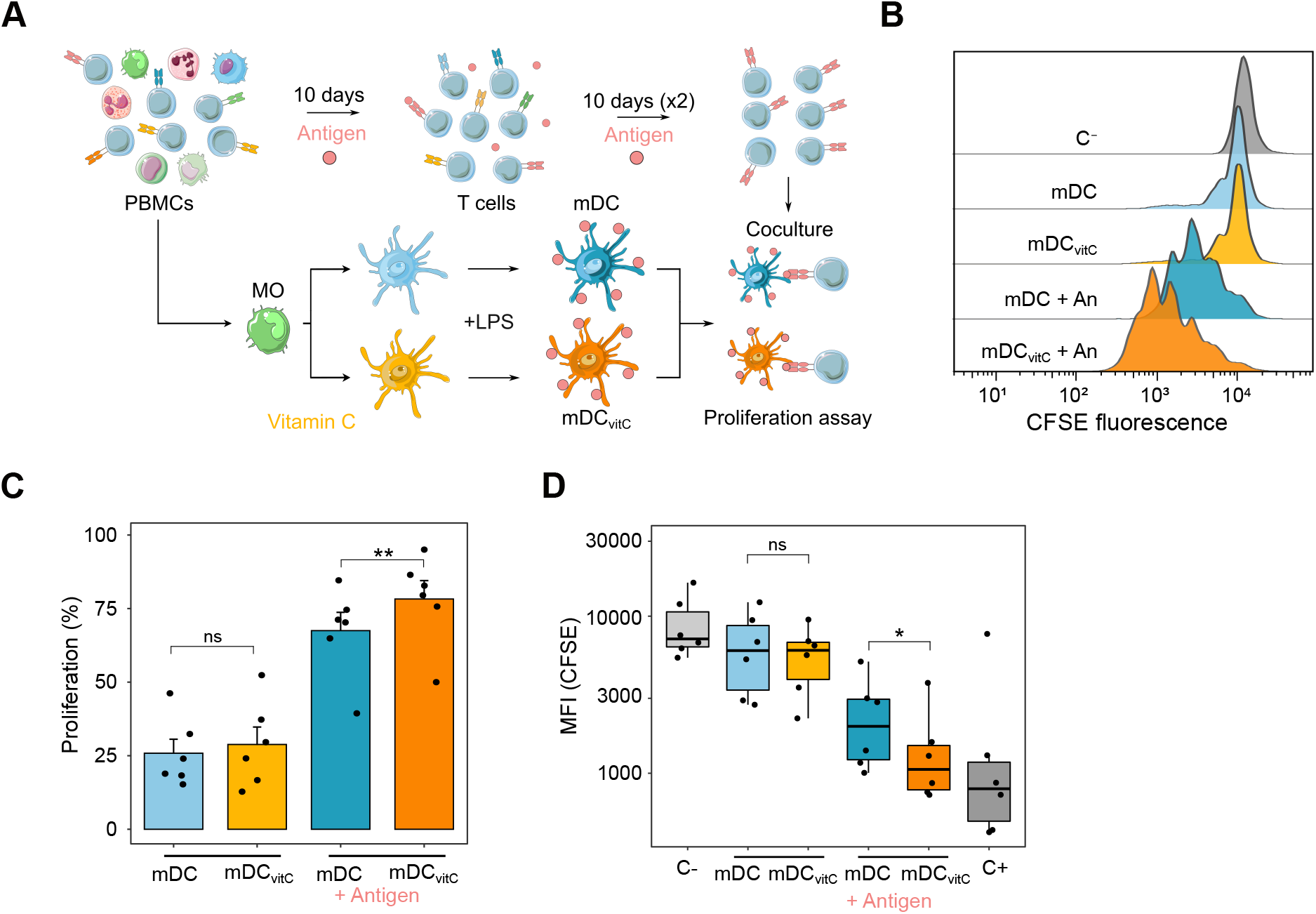
Functional and phenotypic alterations of vitamin C-treated dendritic cells. (A) Scheme depicting the T cell proliferation assay. PBMCs were obtained from healthy donors. T cell clones reacting to the specific antigen (SARS-CoV-2 protein S) were selected through several rounds of clonal expansion. On the other hand, mDCs and mDC_vitC_ were obtained from the same donor and charged with the specific antigen. Finally, the Carboxyfluorescein succinimidyl ester(CFSE)-stained T cells were cocultured with mDCs/mDC_vitC_ for 5 days. (B) Histogram of CFSE signal from T cells, alone without stimulation (C□) or cocultured with mDC/mDC_vitC_ treated with a control antigen or a specific set of antigens of SARS-CoV-2 (An). When T cells proliferate, the CFSE signal is diminished. (C) The proliferation of T cells cocultured with mDC/mDC_vitC_ (2:1 proportion), with or without the loading with a specific set of antigens of SARS-CoV-2. Negative control from each donor was used to calculate the proliferation percentage. P-values from two-tailed paired t-tests are shown (n=6) (ns P > 0.05, ** P < 0.01). (D) Median Fluorescence Intensity (MFI) of T cells alone (C□), cocultured with mDC/mDC_vitC_ (2:1 proportion), with or without the loading with a specific set of antigens of SARS-CoV-2, or stimulated with CD3/CD28 activation beads (C□). P-values from two-tailed paired t-tests are shown (n=6) (ns P > 0.05, * P < 0.05).

In Figure 5B, a selected example of this T cell proliferation assay is shown. The histogram of the CFSE signal of T cells alone (C□) or cocultured with mDC/mDC_vitC_ loaded with a control antigen or with the specific set of antigens (An) is depicted.

Overall, we observed a significant increase in T cell proliferation when they are cocultured with mDC_vitC_ loaded with a specific set of antigens, in comparison with mDCs loaded with the same antigens. This can be observed by the increase in the proliferation percentages and the decrease in the median intensity of fluorescence (MFI) of CFSE (Figure 5C, Figure 5D). However, when T cells were cocultured with mDCs/mDC_vitC_ loaded with a control antigen, no differences in proliferation or MFI were found, indicating that the mDC_vitC_ increased T cell activation capabilities rely on specific antigen presentation and not only indirect stimulation (Figure 5C, Figure 5D).

## DISCUSSION

In this work, we demonstrate a substantial effect of vitamin C supplementation during the MO-to-DC *in vitro* differentiation and maturation. First, we show vast demethylation in DCs treated with vitamin C, consistently with its role as a TET enzyme cofactor, being the number of additional demethylated CpGs higher than the MO-to-DC demethylation without vitamin C. This epigenomic remodeling correlates with increased expression of genes related to antigen presentation, cytokine secretion, and immune response. Moreover, our analysis indicates that NF-kB is directly involved in the epigenomic and transcriptomic reprogramming observed during the maturation of DCs in the presence of vitamin C, together with the increase in TNFβ production. Finally, vitamin C enhances the capacity of DCs to induce the proliferation of autologous T cells through specific antigen presentation.

Vitamin C is a well-established cofactor of TET proteins, with the ability to enhance TET-mediated oxidation of 5-methylcytosine (5mC) into 5-hydroxymethylcytosine (5hmC) and further oxidized methylcytosine derivatives(*26*, *27*). DNA methylation (5mC) is generally considered a repressive epigenetic mark, associated with gene downregulation(*9*, *28*, *29*). Given the absence of proliferation in the MO-to-DC differentiation process(*13*), the observed demethylation should occur through an active mechanism, consistent with previous studies, and be catalyzed by TET, the activity of which is enhanced by vitamin C. Since TET2 is the most expressed TET enzyme in MOs, it is probably driving the active demethylation process, as shown in similar contexts(*7*, *11*, *30*).

Of note, the observed effects of vitamin C on DNA methylation occur in the last days of the differentiation process, since no DNA methylation differences were found with vitamin C treatment on day 2 of differentiation. In contrast, most variance in methylation occurring in the MO-to-iDC transition occurs before day 2 suggesting that, without vitamin C, the function of TET enzymes in this model is progressively diminished. Alternative reducing agents present in the culture medium such as glutathione are less efficient as TET cofactors(*27*, *31*). Thus, the progressive oxidation of these reducing agents may explain the impairment of TET function over time in the absence of vitamin C.

Vitamin C-mediated demethylation occurs during the differentiation (M1 cluster) or the maturation of DCs (M2 cluster). Interestingly, the sets of transcription factor motifs enriched in the M1 and M2 clusters are equivalent to the transcription factors involved in the differentiation and LPS signaling, respectively. This suggests that, overall, vitamin C does not promote the recruitment of TET enzymes through new pathways but boosts the demethylation triggered by preexisting active signaling pathways. This conception is reinforced by the fact that M1 and M2 CpGs are enriched in regions with increasing active chromatin marks (H3K27ac and H3K4me1) in the MO-to-iDC and the iDC-to-mDC transitions, respectively (Supplementary Figure 1D).

The functional relationship between DNA demethylation and gene expression has been extensively studied in several contexts(*7*, *9*, *10*, *12*, *13*). Genes more expressed in mDC_vitC_ than mDC, from E1 and E2 clusters are enriched in M1-associated genes. This establishes a clear temporal relation, suggesting that prior demethylation could be protecting some genes from downregulation during DC maturation. On the other hand, genes from the E5 cluster are enriched in M2-associated genes. In this case, demethylation is occurring at the same step as upregulation. Then, we cannot know if demethylation or upregulation occurs first. However, since the primary mechanism of action of vitamin C through DNA and histones demethylation is well known, we hypothesize that epigenetic modifications could be mediating gene upregulation in mDC_vitC_. This assumption starts from a different point than other works that established DNA demethylation during DC maturation as a consequence of gene upregulation because in that case, the differential stimulus is live bacteria, that can activate a plethora of signaling pathways not necessarily linked directly to DNA demethylation(*10*).

NF-κB mediates both demethylation and upregulation during the maturation of DCs in the presence of vitamin C (Figure 4D, Figure 4E). Since that pathway is associated with tolllike receptor 4 (TLR4), which recognizes LPS, it is not surprising that it plays a role in the process. Intriguingly, this factor is present in the nucleus in mDCs and mDC_vitC_, interacts with TET2, and binds in both cases to CpGs that become demethylated only in mDC_vitC_.

We postulate that NF-kB binding in mDCs and mDC_vitC_ is probably similar, but vitamin C potentiates TET2 function allowing the demethylation of genomic loci, which may condition the expression of the associated genes. Moreover, NF-kB also drives vitamin C-mediated production of TNFβ. This cytokine can signal through tumor necrosis factor receptor (TNFR)I and TNFRII to activate the NF-kB pathway in both DCs and T cells and has shown anti-carcinogenic properties in animal models (32–34).

Linus Pauling proposed vitamin C as a potential cancer treatment more than 40 years ago, but the negative results of further clinical trials diminished the enthusiasm(*35*, *36*). However, during the last few years, increasing interest has arisen around vitamin C as a treatment or adjuvant for several types of cancer. For instance, intravenous vitamin C treatment in mice abrogates cancer progression through direct TET2 function restoration in cancer cells(*16*). Moreover, clinical remission following vitamin C treatment was found in a case of acute myeloid leukemia with mutations in *TET2*(*37*). Furthermore, in mice models of different types of cancer, a fully competent immune system was required to maximize the antiproliferative effects of vitamin C, suggesting an effect of that molecule in the modulation of the immune system(*17*).

On the other hand, immunotherapy with autologous DCs (DC vaccines) has been extensively investigated, with more than 200 completed clinical trials to date(*18*). Most efforts have been focused on cancer, but some clinical trials have also been initiated to treat infectious diseases such as COVID19 (NCT04685603, NCT05007496)(*38*). The use of moDCs differentiated *ex vivo* from monocytes of the same donor is a common and straightforward approach to generating DC vaccines, given the relatively high abundance of these cells in the human blood. However, the lower antigen presentation capabilities of moDCs in comparison with blood DCs is a bottleneck for the efficacy of these treatments(*39*).

Here, we show that mDC_vitC_ loaded with SARS-CoV-2 antigens can stimulate the proliferation of autologous antigen-specific T cells more efficiently than mDCs. This indicates that vitamin C induces an increase of the antigen presentation capabilities of mDCs in an antigen-specific context, and could be a promising strategy for generating DC vaccines towards specific tumor antigens. The improvement in the immune function is linked to epigenomic and transcriptomic remodeling mediated by vitamin C. These results can lead to the generation of new *in vitro* protocols for the generation of moDC vaccines with higher performance. Moreover, it also exposes a potential mechanistic explanation for the vitamin C antineoplastic effects, through *in vivo* modulation of DC differentiation. Further works using animal models and human *in vivo* moDCs from patients treated with high doses of vitamin C should shed light on the specific clinical implications of these insights.

## MATERIALS AND METHODS

### CD14□ monocyte purification and culture

Buffy coats were obtained from anonymous donors via the Catalan Blood and Tissue Bank (CBTB). The CBTB follows the principles of the World Medical Association (WMA) Declaration of Helsinki. Before providing blood samples, all donors received detailed oral and written information and signed a consent form at the CBTB.

PBMCs were isolated by density-gradient centrifugation using lymphocyte-isolation solution (Rafer). Pure MOs were then isolated from PBMCs by positive selection with magnetic CD14 MicroBeads (Miltenyi Biotec). Purity was verified by flow cytometry, which yielded more than 95% of CD14□ cells.

MOs were resuspended in Roswell Park Memorial Institute (RPMI) Medium 1640 + GlutaMAX™ (Gibco, ThermoFisher) and immediately added to cell culture plates. After 20 minutes, monocytes were attached to the cell culture plates, and the medium was changed with RPMI containing 10% fetal bovine serum (Gibco, ThermoFisher), 100 units/mL penicillin/streptomycin (Gibco, ThermoFisher), 10 ng/mL human GM-CSF (PeproTech) and 10 ng/mL human IL-4 (PeproTech). In the case of the cells treated with vitamin C, 500 μM (+)-Sodium L-ascorbate (Sigma-Aldrich) was also added to the medium. For dendritic cell maturation, LPS (10 ng/mL) was added to cell culture on day 5.

Cells were collected on day 2 (Ø / Ø_vitC_) and on day 7, including immature dendritic cells (iDC / iDC_vitC_) and mature dendritic cells, with LPS stimulus (mDC / mDC_vitC_).

### Genomic DNA and total RNA extraction

Genomic DNA and total RNA were extracted using the Maxwell RSC Cultured Cells DNA kit (Promega) and the Maxwell RSC simplyRNA Cells kit (Promega), respectively, following the manufacturer’s instructions.

### DNA methylation profiling

500 ng of genomic DNA was converted using the EZ DNA Methylation Gold kit (Zymo Research), using 4 biological replicates for each group. Infinium MethylationEPIC BeadChip (Illumina) arrays were used to analyze DNA methylation, following the manufacturer’s instructions. This platform allows around 850,000 methylation sites per sample to be interrogated at single-nucleotide resolution, covering 99% of the reference sequence (RefSeq) genes. Raw files (IDAT files) were provided for the Josep Carreras Research Institute Genomics Platform (Barcelona).

Quality control and analysis of EPIC arrays were performed using ShinyÉPICo(*40*) a graphical pipeline that uses minfi(*41*) for normalization, and limma(*42*) for differentially methylated positions analysis. CpH and SNP loci were removed and the Noob+Quantile normalization method was used. Donor information was used as a covariate, and Trend and Robust options were enabled for the eBayes moderated t-test analysis. CpGs were considered differentially methylated when the absolute differential of methylation was greater than 20% and the FDR was less than 0.05.

### RNA-seq

RNA-seq libraries of MOs, Ø / Ø_vitC_, iDC / iDC_vitC_, and mDC / mDC_vitC_ were generated and sequenced by Novogene (Cambridge), in 150-bp paired-end, with the Illumina NovaSeq 6000 platform, using 3 biological replicates for each group. More than 40 million reads were obtained for each sample. Fastq files were aligned to the hg38 transcriptome using HISAT2(*43*) with standard options. Reads mapped in proper pair and primary alignments were selected with SAMtools(*44*). Reads were assigned to genes with featureCounts(*45*).

Differentially expressed genes were detected with DESeq2 (*47*). The donor was used as a covariate in the model. The Ashr shrinkage algorithm was applied and only proteincoding genes with an absolute logFC greater than 0.5 and an FDR less than 0.05 were selected as differentially expressed. For representation purposes, Variance Stabilizing Transformation (VST) values and normalized counts provided by DESeq2 were used.

### NF-κB chemical inhibition

Bay 11-7082 (Sigma-Aldrich) was used for NF-κB inhibition. The compound was diluted in DMSO to 50mM. Bay 11-7082 at 10μM or an equivalent amount of diluent were used as final concentrations.

MOs were differentiated to iDCs/iDC_vitC_ as previously described. On day 5, LPS, Bay 11-7082 or equivalent amounts of diluents were added to the cell culture for 2 days, yielding iDC/iDC_vitC_, iDC+Bay/iDC_vitC_+Bay, mDC/mDC_vitC_, and mDC + Bay/mDC_vitC_ + Bay.

### Quantification of cytokine production

Cell culture supernatants were collected after 7 days and diluted appropriately. Enzyme-linked immunosorbent assays (ELISA) were performed to detect TNFβ, following the manufacturer’s instructions (TNF beta Human ELISA Kit, ThermoFisher).

### T cell clonal expansion and proliferation assay

PBMCs from healthy donors were purified from blood by density gradient centrifugation. The PBMCs (1ml; 3·10^6^ cells per well) were cultured in the presence of SARS-CoV-2-S (9pmol) (PepTivator) in 24-well plates and maintained in IMDM medium (Gibco) supplemented with penicillin (100 units/mL), streptomycin (100 mg/mL) and human serum (10%) (Millipore) in the absence of IL-2 for 3 days. After 3 days, 1ml of medium with 80 U/mL of recombinant human IL-2 (PeproTech) was added to the wells, with a final concentration of 1.5·10^6^ cell/ml and 40U/ml of IL-2. After 7-10 days of culture, T cells were expanded in the presence of 30-Gy irradiated autologous PBMCs (3·10^6^ cells/well) previously pulsed with 9pmol of SARS-CoV-2-S (PepTivator). Antigen-specific T cells (mix of CD4+ and CD8+ T cells) were selected by performing the same protocols two times to have a positive selection.

After 7 days of differentiation and activation, DCs were washed to remove vitamin C and were co-cultured with antigen-specific autologous CFSE-stained T cells at a DC: T cell ratio of 1:2 in 200μl of RPMI 1640 medium containing 10% FBS, penicillin (100 units/mL), streptomycin (100 mg/mL) in round bottom 96-well plates (ThermoFisher). Co-culture was performed in the presence of SARS-CoV-2-S antigen or SARS-CoV-2-N control antigen (PepTivator). T cell proliferation was analyzed by FACS and determined by considering the proliferating of those where CFSE staining had decreased compared to not co-cultured T cells. T cells stimulated with anti-CD3/CD28 microbeads 5ug/mL (eBioscience) were used as a positive control.

### Flow cytometry

To study cell-surface markers, cells were collected using Versene, a non-enzymatic dissociation buffer (ThermoFisher). Cells were resuspended in the staining buffer (PBS with 4% fetal bovine serum and 2 mM ethylenediaminetetraacetic acid (EDTA)). Cells were then incubated on ice with an Fc block reagent (Miltenyi Biotec) for 10 minutes, and stained with the proteins of interest, using the following antibodies: CD8 (FITC) (#21270083, Immunotools), and CD4 (APC) (#555349, BD Biosciences). After staining, cells were analyzed using a BD FACSCanto™ II Cell Analyzer (BD Biosciences). Data were analyzed with the FlowJo v10 software.

### Bisulfite pyrosequencing

500 ng of genomic DNA was converted using the EZ DNA Methylation Gold kit (Zymo Research). PCR was performed using the bisulfite-converted DNA as input and primers designed for each amplicon (Supplementary Table 4). These primers were designed using the PyroMark Assay Design 2.0 software (Qiagen). PCR amplicons were pyrosequenced using the PyroMark Q48 system and analyzed with PyroMark Q48 Autoprep software.

### Real-time quantitative reverse-transcribed polymerase chain reaction (qRT-PCR)

300 ng of total RNA were reverse-transcribed to cDNA with Transcriptor First Strand cDNA Synthesis Kit (Roche) following the manufacturer’s instructions. qRT-PCR was performed in technical triplicates for each biological replicate, using LightCycler^®^ 480 SYBR Green Mix (Roche), and 7.5 ng of cDNA per reaction. The average value from each technical replicate was obtained. Then, the standard double-delta Ct method was used to determine the relative quantities of target genes, and values were normalized against the control genes RPL38 and HPRT1. Custom primers were designed to analyze genes of interest (Supplementary Table 4).

### Co-immunoprecipitation (Co-IP)

Co-IP assays were performed using mDCs and mDC_vitC_ after 5 days of differentiation from MOs and 24h of stimulation with LPS. Cell extracts were prepared in lysis buffer [50 mM Tris–HCl, pH 7.5, 1 mM EDTA, 150 mM NaCl, 1% Triton-X-100, protease inhibitor cocktail (cOmplete™, Merck)] with corresponding units of Benzonase (Sigma) and incubated at 4°C for 4 h. 100 μl of supernatant was saved as input and diluted with 2× Laemmli sample buffer (5x SDS, 20% glycerol, 1M Tris–HCl (pH 8.1)). Supernatants were first precleared with PureProteome™ Protein A/G agarose suspension (Merck Millipore) for 1 h. The lysate was then incubated overnight at 4°C with respective crosslinked primary antibodies. The cross-linking was performed in 20mM dimethyl pimelimidate (DMP) (Pierce, Thermo Fisher Scientific, MA, USA) dissolved in 0.2 M sodium borate (pH 9.0). Subsequently, the beads were quenched with 0.2M of ethanolamine (pH 8.0) and resuspended at 4°C in PBS until use. Beads were then washed three times with lysis buffer at 4°C. Sample elution was done by acidification using a buffer containing 0.2 M glycine (pH 2.3) and diluted with 2× Laemmli. Samples and inputs were denatured at 95°C in the presence of 1%β-mercaptoethanol. Antip65 C15310256 (DIagenode) and control IgG C15410206 (Diagenode) were used for Co-IP.

### Chromatin Immunoprecipitation

On day 6, after 5 days of differentiation and 24h of maturation with LPS or an equivalent amount of diluent, iDC, iDC_vitC_, mDC, and mDC_vitC_ were fixed with Pierce™ fresh methanol-free formaldehyde (ThermoFisher) for 15min and prepared for sonication with the truChIP Chromatin Shearing Kit (Covaris), following the manufacturer’s instructions. Chromatin was sonicated 18min with the Covaris M220 in 1mL milliTubes (Covaris). The size distribution of the sonicated chromatin was checked by electrophoresis to ensure an appropriate sonication, with a size around 200bp.

Magna Beads Protein A+G (Millipore) were blocked with PBS + BSA (5mg/mL) for 1hour. Chromatin was precleared with 25μl of beads for 1.5h and 10μg of chromatin were incubated overnight with each antibody: 10μl Anti-p65 antibody ab16502 (Abcam), in a buffer with 1% Triton X-100, 150mM NaCl, and 0.15% SDS. Then, 3 washes were performed with the Low Salt Wash Buffer (0.1% SDS, 1%Triton X-100, 2mM EDTA, pH 8.0, 20mM Tris-HCl pH 8.0, 150mM NaCl), the High Salt Wash Buffer (0.1%SDS, 1% Triton X100, 2mM EDTA, pH 8.0, 20mM Tris-HCl pH 8, 500mM NaCl), and the LiCl Wash Buffer (0.25 M LiCl, 1% Nonidet P-40, 1% Deoxycholate, 1mM EDTA pH 8, 10mM Tris-HCl), followed by a last wash with TE buffer (pH 8.0, 10mM Tris-HCl, 1mM EDTA). Chromatin was eluted for 45min 65°C with 100μl of elution buffer (10mM Tris-Cl, 1mM EDTA, 1%SDS) and decrosslinked adding 5μl 5M NaCl and 5μl 1M NaHCO3 (2h 65°C). Next, 1μl of 10mg/mL proteinase K (Invitrogen) was added, and samples were incubated at 37°C for 1h. For DNA purification, iPure kit v2 (Diagenode) was used, following the manufacturer’s instructions. 1% of the chromatin input from each sample was purified by the same method.

For ChIP-qPCR, samples were diluted 1/10, and 4μL and specific primers (Supplementary Table 4) were used for each reaction. qRT-PCR was performed in technical triplicates for each biological replicate, using LightCycler^®^ 480 SYBR Green Mix (Roche). The relative amount of immunoprecipitated DNA compared to input was calculated with the following formula: 2 ^((Ct_input_ – 6.64) - Ct_sample_) * 100%

### Western Blotting

Cytoplasmic and nuclear protein fractions were obtained using hypotonic lysis buffer (Buffer A; 10 mM Tris pH 7.9, 1.5 mM MgCl_2_, 10 mM KCl supplemented with protease inhibitor cocktail (Complete, Roche) and phosphatase inhibitor cocktail (PhosSTOP, Roche) to lyse the plasma membrane. Cells were visualized in the microscope to assure correct cell lysis. The nuclear pellets were resuspended in Laemmli 1X loading buffer. For whole-cell protein extract, cell pellets were directly resuspended in Laemmli 1X loading buffer.

Proteins were separated by SDS-PAGE electrophoresis. Immunoblotting was performed on polyvinylidene difluoride (PVDF) membranes following standard procedures. Membranes were blocked with 5% Difco™ Skim Milk (BD Biosciences) and blotted with primary antibodies. After overnight incubation, membranes were washed three times for 10 minutes with TBS-T (50 mM Tris, 150 mM NaCl, 0.1% Tween-20) and incubated for 1 hour with HRP-conjugated mouse or rabbit secondary antibody solutions (Thermo Fisher) diluted in 5% milk (diluted 1/10000). Finally, proteins were detected by chemiluminescence using WesternBright™ ECL (Advansta). The following antibodies were used: Anti-p65 C15310256 (DIagenode), Anti-phosphorylated p65 (Ser536) 93H1 (Cell Signaling), Anti-GAPDH 2275-PC-100 (Trevigen), Anti-TET2 C15200179 (Diagenode), Anti-histone H1 ab4269 (Abcam), Anti-beta Actin ab8227 (Abcam). Protein quantification was performed with ImageJ/Fiji (*46*).

### Data analysis and representation

Statistical analyses were performed in R 4.0.3. Gene expression and DNA methylation heatmaps were created with the heatmap.2 function of the gplots package. The findMotifsGenome.pl function of HOMER (Hypergeometric Optimization of Motif EnRichment) was used to analyze known motif enrichment, using the parameters ‘-size 200 -cpg. All EPIC array CpG coordinates were also used as background for the methylation data. GREAT software was used to calculate CpG-associated genes and gene ontology (GO) enrichment(*47*). GO enrichment of gene expression data was performed using the clusterProfiler package(*48*). ChIP-seq peaks files of histone marks from MO, iDCs, and mDCs were downloaded from the BLUEprint webpage (http://dcc.blueprint-epigenome.eu). Consensus peaks of the different replicates were obtained with the MSPC algorithm(*49*), using the options ‘-r Biological -w 1E-4 -s 1E-8 -c 3’.

The chromatin state learning model for CD14+ monocytes was downloaded from the Roadmap Epigenomics Project webpage, and chromatin state enrichments were calculated using Fisher’s exact test.

Proportional Venn diagrams were generated with the Meta-Chart webpage (https://www.meta-chart.com/).

## DATA AVAILABILITY

DNA methylation and expression data for this publication have been deposited in the NCBI Gene Expression Omnibus and are accessible through GEO SuperSeries accession number GSE203463.

## ACKNOWLEDGEMENTS

We thank CERCA Programme/Generalitat de Catalunya and the Josep Carreras Foundation for institutional support. E.B. was funded by the Spanish Ministry of Science and Innovation (PID2020-117212RB-I00; AEI/10.13039/501100011033). J.L.S. was funded by Instituto de Salud Carlos III through the project CP19/00176 (co-funded by European Social Fund, “Investing in your future”) and the Spanish Ministry of Science, Innovation and Universities (MICINN; grant number PID2019-111243RA-I00 / AEI / 10.13039/501100011033). E.M.C. is funded by project PI20/01313, integrated in the Plan Nacional de I + D + I and co-supported by the ISCIII-Subdirección General de Evaluación and the Fondo Europeo de Desarrollo Regional (FEDER). O.M.-P. holds an i-PFIS PhD fellowship (grant number IFI17/00034) from Acción Estratégica en Salud 2013-2016 ISCIII, co-financed by Fondo Social Europeo.

## AUTHOR CONTRIBUTIONS

O.M.-P. and E.B. conceived and designed the study; O.M.-P, G.G.-T., J.C.-S., L.C., E.M.-C. and J.L.S. performed experiments; O.M.-P performed the bioinformatic analyses; O.M.-P and E.B. wrote the manuscript; all authors participated in discussions and interpreting the results.

