## Supplementary figures and images for "Vitamin C triggers NF-κB-driven epigenomic reprogramming and enhanced immunogenic responses of dendritic cells"

### Supplementary Figure 1

#### Supplementary Figure 1

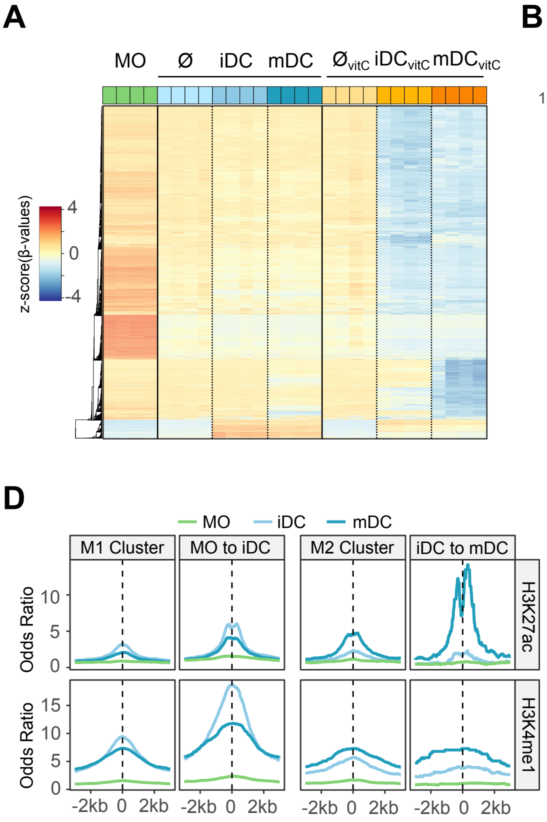

### Supplementary Figure 2

#### Supplementary Figure 2

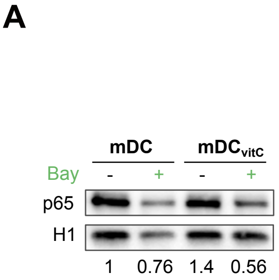


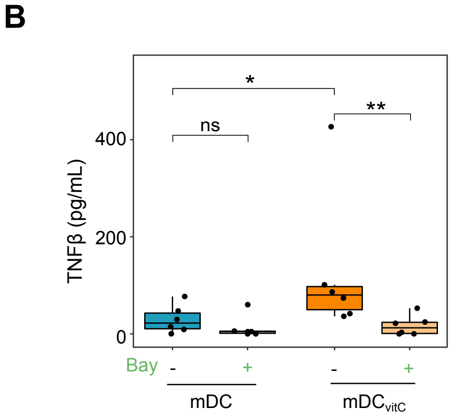

### Supplementary Figure 3

#### Supplementary Figure 3

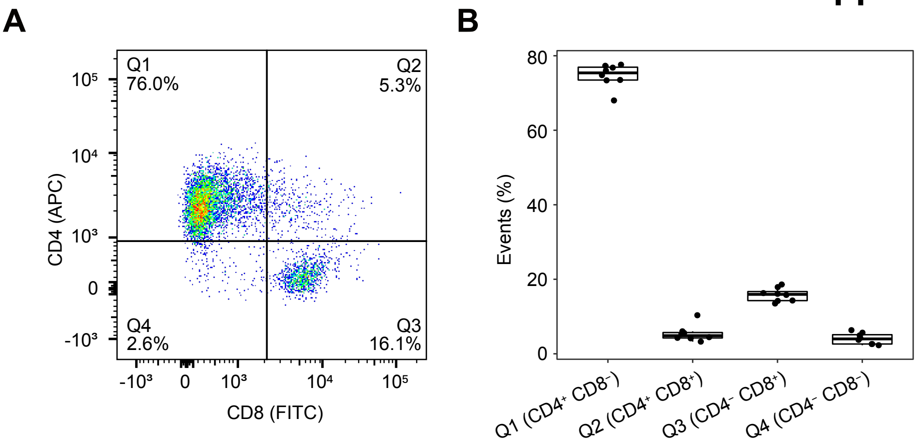
