## Supplementary Figure Legends for "Vitamin C triggers NF-κB-driven epigenomic reprogramming and enhanced immunogenic responses of dendritic cells"

**Supplementary Figure 1.** (A) DNA methylation heatmap of differentially methylated CpGs between all groups, compared pairwise ($\Delta\beta$- ≥ 0.3, FDR < 0.05). Scaled β-values are shown (lower DNA methylation levels in blue and higher methylation levels in red). (B) Barplot of genomic features percentages of M1 and M2 CpGs in comparison with background CpGs (Bg). (C) Barplot of CpG island contexts percentages of M1 and M2 CpGs in comparison with background CpGs. (D) ChIP-seq data of H3K27ac and H3K4me1 of CD14+ MOs, iDCs, and mDCs were downloaded from the BLUEPRINT database. Odds ratios were calculated for bins of 10 bp up to 2000 bp around M1 and M2 CpGs. CpGs annotated in the EPIC array were used as background.

**Supplementary Figure 2.** (A) Western blot of p65 in the nuclear fraction of mDC and $mDC_{vitC}$. Histone H1 was used as a loading control. Signal of western blot bands was quantified and p65/H1 ratios are shown below each sample. (B) Boxplot depicting the effect of NF-κB inhibition with BAY 11-7082 (Bay) in the production of TNFβ by mDCs and $mDC_{vitC}$, as quantified in cell supernatants. The P-value of a Wilcoxon rank-sum test is shown (n=6) (ns $P>0.05$, *$P < 0.05$, **$P < 0.01$).

**Supplementary Figure 3.** (A) Example of T cell staining after the three rounds of clonal expansion. The fluorescence of CD4 is shown on the y-axis, whereas the fluorescence of CD8 is shown on the x-axis. The four quadrants are divided depending if the cell is positive for CD4 (Q1), CD8 (Q3), both (Q2), or none (Q4). (B) Boxplot showing the percentages of each quadrant from the CD4/CD8 staining (n=8).
